# Asymmetric wall ingrowth deposition in Arabidopsis phloem parenchyma transfer cells is tightly associated with sieve elements

**DOI:** 10.1101/2022.01.04.475004

**Authors:** Xiaoyang Wei, Yuan Huang, David A. Collings, David W. McCurdy

## Abstract

In Arabidopsis, polarized deposition of wall ingrowths in phloem parenchyma (PP) transfer cells (TCs) occurs adjacent to cells of the sieve element/companion cell (SE/CC) complex. However, the spatial relationships between these different cell types in minor veins, where phloem loading occurs, are poorly understood. PP TC development and wall ingrowth localization were compared to other phloem cells in leaves of Col-0 and the transgenic lines *AtSUC2::AtSTP9-GFP* and *AtSWEET11::AtSWEET11-GFP* that identify CCs and PP respectively. The development of PP TCs in minor veins, indicated by deposition of wall ingrowths, proceeded basipetally in leaves. However, not all PP develop ingrowths and higher levels of wall ingrowth deposition occur in abaxial-compared to adaxial-positioned PP TCs. Furthermore, the deposition of wall ingrowths was exclusively initiated on and preferentially covered the PP TC/SE interface, rather than the PP TC/CC interface, and only occurred in PP cells that were adjacent to SEs. Collectively, these results demonstrate the dominant impact of SEs on wall ingrowth deposition in PP TCs and suggest the existence of two sub-types of PP cells in leaf minor veins. Compared to PP cells, PP TCs showed more abundant accumulation of AtSWEET11-GFP, indicating functional differences in phloem loading between PP and PP TCs.

**Highlight:** Wall ingrowth deposition in phloem parenchyma transfer cells of Arabidopsis leaf minor veins is initiated adjacent to sieve elements, rather than companion cells, and expands to preferentially cover this interface.

## Introduction

The phloem is the main long-distance transport system for nutrients in plants. Anatomically, phloem is built up by arrays of companion cells (CCs), sieve elements (SEs), and adherent phloem parenchyma (PP) cells, in which CCs and SEs are connected by plasmodesmata and thus form the SE/CC complex which acts as the fundamental structural and functional unit of the phloem system (van Bel, 2003; Turgeon and Wolf, 2009). In some species such as pea and *Arabidopsis thaliana* (Arabidopsis), phloem cells, including both CCs and PP cells, deposit extensive networks of wall ingrowths, *trans*-differentiating into transfer cells (TCs) (Pate and Gunning, 1969; Amiard et al., 2007; Nguyen and McCurdy, 2016). In Arabidopsis, TCs *trans*-differentiate from PP and occur exclusively in the minor veins of foliar tissues such as cotyledons and leaves (Haritatos et al., 2000; Nguyen and McCurdy, 2015). A survey of this process revealed that wall ingrowth deposition in these minor veins is a novel trait of heteroblasty, being more abundant in fully mature juvenile leaves compared to mature adult leaves. In addition, the level of wall ingrowth deposition displays a distinct basipetal gradient in mature adult leaves with ingrowths being more developed in the leaf apices (Nguyen et al., 2017). However, the fine-scale developmental progression of PP TCs across leaf development remains unclear.

TCs undergo wall ingrowth deposition to increase plasma membrane surface area thus facilitating increased capacity for trans-membrane transport. In Arabidopsis, PP TCs in foliar minor veins play important roles in phloem loading by importing sucrose symplasmically from bundle sheath cells then exporting this sucrose into the apoplasm for subsequent uptake by CCs in the SE/CC complex (Chen, 2014). Consistent with their proposed role in phloem loading, wall ingrowth deposition in PP TCs occurs exclusively along the interface adjacent to the SE/CC complex (Haritatos et al., 2000; Amiard et al., 2007; Nguyen et al., 2017), which is distinct from CC TCs in which wall ingrowth deposition occurs uniformly along the interface of the primary cell wall (Pate and Gunning, 1969). This polarized distribution of wall ingrowth deposition in PP TCs, in turn, implies potential influence of the SE/CC complex on the formation of these structures in PP TCs. However, observations in the literature suggest that wall ingrowth deposition is not only associated with CCs but also associated with SEs (Haritatos et al., 2000; Amiard et al., 2007; Nguyen et al., 2017). Nonetheless, the exact distribution of wall ingrowth deposition in the PP TCs, and precise relationships with the abutting SE/CC complex, has not been fully elucidated.

Recent research has demonstrated additional complexity within the organization of vascular bundles within leaves. Chen et al. (2018) identified two sub-types of CCs in minor veins of Arabidopsis, either those that express *FLOWERING LOCUS T (FT)* or those that do not. Similarly, transcriptomes of abaxial and adaxial bundle sheath cells in maize leaf minor veins are distinct from each other (Bezrutczyk et al., 2021). Furthermore, recent single-cell RNA-sequencing (scRNA-Seq) of enriched leaf vascular tissue reported the identification of two clusters of PP cells, one enriched in defence and cell wall genes (PP1 cluster) and one enriched in photosynthesis-related genes (PP2 cluster) (Kim et al., 2021). Consequently, one possibility arising from this study is that not all PP cells in leaf minor veins *trans*-differentiate to become PP TCs by deposition of wall ingrowths. Since both AtSWEET11 and AtSWEET12 are cell-type markers of PP cells in foliar tissues in Arabidopsis (Chen et al., 2012; Cayla et al., 2019), a structural survey of transgenic lines expressing *AtSWEET11-GFP* or *AtSWEET12-GFP* constructs may answer this question.

In this study, the developmental processes and morphology of PP TCs and their spatial relation with other phloem cells were surveyed in Arabidopsis leaves. To assess cellular organisation in minor veins, vibratome sectioning allowing high spatial resolution of vein structure was undertaken. Additionally, structural studies were conducted on leaf minor veins in transgenic fluorescent plants expressing the pAtSUC2::AtSTP9-GFP and *p*AtSWEET11::AtSWEET11-GFP constructs that identify different phloem cell types. Observation and quantification of these lines was possible using the ClearSee imaging technique, which allowed observation of the fluorescent proteins concurrently with stained wall ingrowths. In leaves, the development of PP TCs coincided with the onset of phloem loading in the plant. In addition, accumulation of AtSWEET11 positively correlated with wall ingrowths in PP cells. These observations indicate the relation between wall ingrowth deposition and the phloem loading activity in PP TCs is interactive. However, tight association between wall ingrowth deposition and SEs was observed.

## Materials and methods

### Plant growth and materials

Seed of plants expressing *pAtSUC2::AtSTP9-GFP* (*tmSTP9*) and *pAtSWEET11::AtSWEET11-GFP were* kindly provided by Dr. Ruth Stadler (Universität Erlangen-Nürnberg, Germany) and Dr. Sylvie Dinant (INRAE, France), respectively, while Col-0 seeds were supplied by the ABRC. All seeds were sown and germinated in potting mix soil after being stratified in darkness at 4°C for 48 h. After stratification, all plants were grown in a growth cabinet under standard lighting conditions for Arabidopsis (100–120 μmol.m^−2^.sec^−1^, 22°C day/18°C night, 16 h photoperiod).

### Cross sectioning and ClearSee treatment

Transition leaves from 4- to 5-week-old seedlings (precise leaf number and age of seedlings sampled in each experiment are stated in the corresponding Results section) of Col-0 and *pAtSUC2::AtSTP9-GFP* were fixed in 4% (w/v) formaldehyde in phosphate buffered saline (PBS: 137 mM NaCl, 2.7 mM KCl, 10 mM Na_2_HPO_4_, 1.8 mM KH_2_PO_4_) in a vacuum chamber at room temperature for 1 h. Fixed tissues were then washed thoroughly in PBS before being embedded in 4% (w/v) agarose gel blocks prepared in TAE buffer (40 mM Tris-acetate, 1 mM EDTA, pH 8.0). Cross sections were cut with a Leica VT1200 vibratome, with the section thickness of 150 μm.

Mature juvenile leaves 1 and 2 (hereafter described as leaf 1) from 27-day-old *pAtSWEET11::AtSWEET11-GFP* seedlings were sampled for ClearSee treatment. The abaxial epidermis of leaf samples was peeled off as described by Cayla et al. (2019). Thereafter, processed samples were fixed with 4% (w/v) formaldehyde in PBS in a vacuum chamber at room temperature for 1 h, washed with PBS for 1 min twice, then cleared in ClearSee solution (0.66 M xylitol, 0.38 M sodium deoxycholate, 4.16 M urea) for at least four days as described by Kurihara et al. (2015).

### Cell wall labelling and confocal imaging

Propidium iodide (PI) staining of leaf tissues was performed as described by Nguyen and McCurdy (2015). For *pAtSWEET11::AtSWEET11-GFP* plants, cleared samples were stained with 0.05% (w/v) calcofluor white solution for 40 min. After staining, samples were washed in ClearSee solution for 30 min before being mounted on slides in ClearSee for confocal microscopy observation.

Three confocal microscopy systems were used for image collecting in this study, namely the Olympus FV1000 and FV3000 systems and a Leica SP8 system. Imaging settings were as follows: PI, excitation: 552 nm, emission window: 570 to 670 nm; calcofluor white, excitation: 408 nm; emission window: 425 to 460 nm. GFP, excitation: 488 nm, emission window: 497 to 527 nm.

### Measurements of cell/cell interface length and fluorescence intensity

Lengths of the PP TC/CC and PP TC/SE interfaces and the relative fluorescence intensity of SWEET11-GFP were measured using the image processing software ImageJ (Fiji, 1.51n). For the fluorescence intensity assay of SWEET11-GFP, in each comparison set, confocal images from the same sequential scanning were collected for analysis. Confocal images were uploaded into ImageJ and converted into 8-bit images, and the threshold of each image was adjusted using the default setting of the software. Thereafter, fluorescence intensity of the selected region of interest (ROI) was measured. In each minor vein, the relative fluorescence intensity of SWEET11-GFP in PP relative to PP TCs was calculated using the formula:

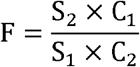

where F is the relative fluorescence intensity. Fluorescence intensity reads in each comparison were as follows:

S_1_, SWEET11-GFP in PP TC;
S_2_, SWEET11-GFP in PP;
C_1_, Calcofluor white staining in PP TC;
C_2_, Calcofluor white staining in PP.

The relative fluorescence intensity of SWEET11-GFP in PP TC was normalized to 1.

In a PP TC, the relative fluorescence intensity of SWEET11-GFP at sites with no wall ingrowths relative to sites containing wall ingrowth deposition was calculated using the formula described above. Fluorescence intensity reads in each comparison were:

S_1_, SWEET11-GFP at the site with wall ingrowths;
S_2_, SWEET11-GFP at the site with no wall ingrowths;
C_1_, Calcofluor white staining at the site with wall ingrowths;
C_2_, Calcofluor white staining at the site with no wall ingrowths.

The relative fluorescence intensity of SWEET11-GFP at the site with wall ingrowths was normalized to1.

### Measurements of plasma membrane enrichment

The ratio of plasma membrane areas comparing sites with and without wall ingrowth deposition was determined by measuring the accumulation of SWEET11-GFP fluorescence. Confocal images were processed in ImageJ as described in Section 3.2.4. For each PP TC, the fold change in plasma membrane area resulting from wall ingrowth deposition was calculated using the formula: where FC in the relative enrichment of plasma membrane area associated with the wall ingrowth relative to the side of the PP TC with no wall ingrowths. Reads of the accumulation area of AtSWEETll-GFP and primary cell wall length in ROI were as follows:

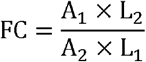

where FC in the relative enrichment of plasma membrane area associated with the wall ingrowth relative to the side of the PP TC with no wall ingrowths. Reads of the accumulation area of AtSWEET11-GFP and primary cell wall length in ROI were as follows:

A_1_, The sum of SWEET11-GFP fluorescence at the site with wall ingrowths;
A_2_: The sum of SWEET11-GFP at the site with no wall ingrowths;
L_1_: Primary cell wall length at the site with wall ingrowths;
L_2_: Primary cell wall length at the site with no wall ingrowths.

Thus, this equation measures the relative fluorescence intensity per unit length of cell wall. These measurements were then compared to the wall ingrowth score in each corresponding PP TC.

## RESULTS

### Basipetal progression of PP TC development in Arabidopsis leaves

Based on the order of their formation and position, veins in Arabidopsis leaves can be classified into five orders (Kang et al., 2007) as indicated in Fig. 1A. Although a previous study indicated the heteroblastic development of PP TCs in Arabidopsis (Nguyen et al., 2017), the fine-scale development of PP TCs along all veins of different orders in different foliar tissues has not been documented. To elucidate this information, the extent of PP TC development and wall ingrowth deposition in veins of each order in developing leaves were scored using the method of Nguyen et al. (2017).

**Fig. 1.**
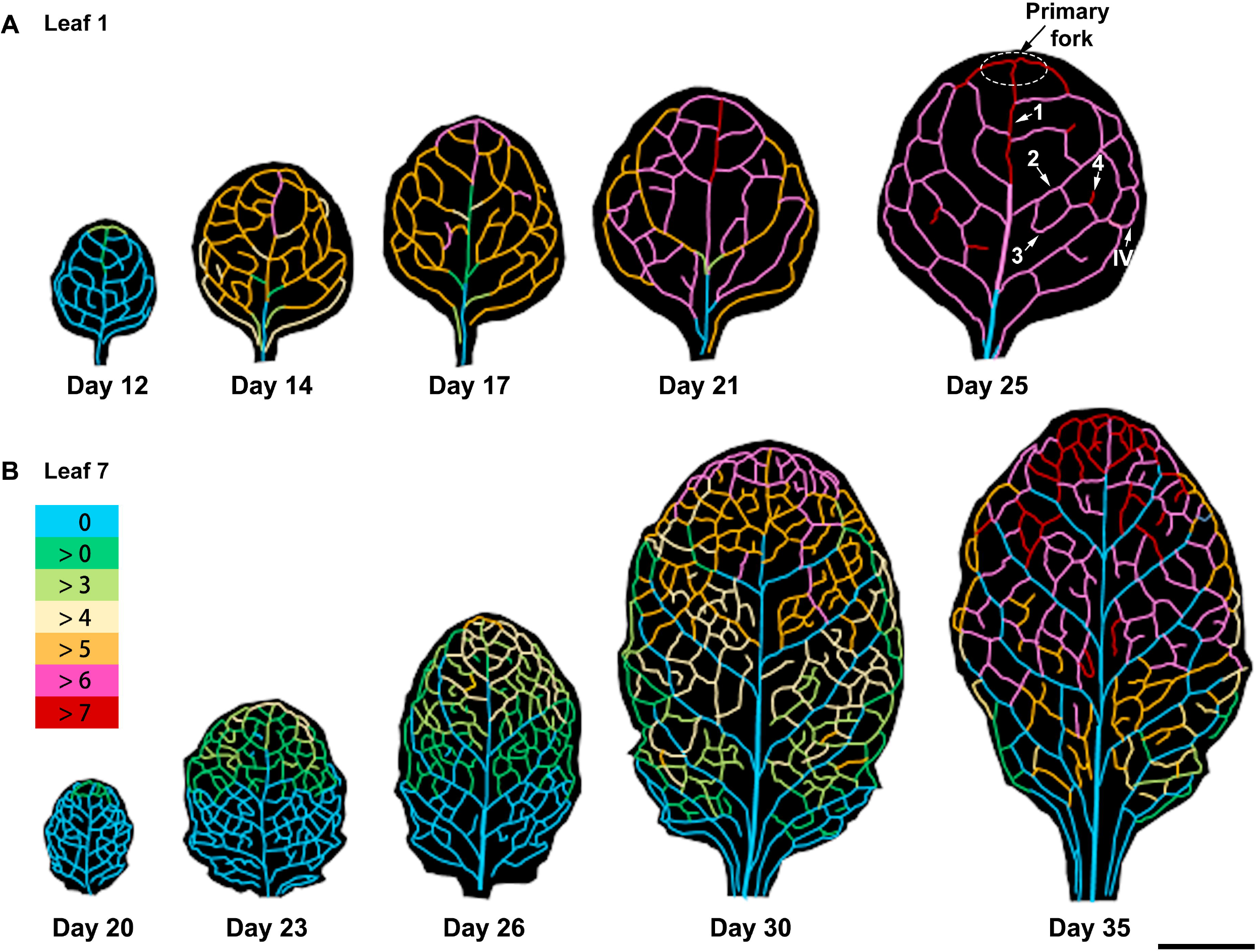
Schematic illustration of PP TC development in Arabidopsis leaves. Wall ingrowths in soil-grown Col-0 plants were quantified following Nguyen et al. (2017) in different sections of veins of each order. The shapes and vein patterns of the representative images are based on leaf scans, with the different colours representing wall ingrowth deposition scores derived from 3 to 7 leaves at each growth stage, and from 1 to 4 confocal images per vein. **(A)** Juvenile leaves (leaf 1) from 12- to 25-day-old plants. **(B)** Leaf 7 from 20- to 35-day-old plants. Vein classes are indicated by arrows in leaf 1 of a 25-day-old plant: (**1**) Midrib, (**2**) secondary vein, (**3**) tertiary vein, (**4**) quaternary vein, and (IV) intramarginal vein. The primary fork is circled and indicated by an arrow. The key in **B** shows the colours denoting PP TC score. Scale bar: 0.5 cm.

Wall ingrowth abundance in PP TCs followed a distinct basipetal gradient in mature leaves including leaf 7 (Fig. 1B). Ingrowth deposition was abundant in the apical ends of the midrib and the secondary veins (indicated by 1 and 2, respectively), but was not observed in the remainder of these veins (Fig. 1B). Furthermore, veins of the same order that were close to the leaf apex had higher levels of ingrowth deposition compared to veins located in the leaf base (Fig. 1B). In leaf 1, however, this developmental gradient was only observed in the midrib and secondary veins, with only the midrib having a strong basipetal gradient (Fig. 1A; Supplementary Fig. 1). In addition, in mature leaves, lower order veins (indicated by 3 and 4 in Fig. 1A), formed later and displayed more abundant wall ingrowths compared to higher order veins (indicated by 1 and 2 in Fig. 1A).

How is this basipetal distribution pattern formed? One possibility is that wall ingrowth accumulation in the leaf apex and the base were initiated at the same time but occurred faster in the apical sites compared to the basal sites. Another possibility is that the onset of wall ingrowth deposition occurred earlier in the leaf apex than the leaf base. A time-course survey supported the second possibility, and that the process of wall ingrowth accumulation in leaves proceeded basipetally (Fig. 1). The onset of wall ingrowth deposition first occurred at the leaf apex and then gradually proceeded to the leaf base (Fig. 1). Consequently, the veins of the primary fork (the first vein that forms branching from the midrib) started to form ingrowths earlier than other veins, and thus deposited more abundant wall ingrowths (Fig. 1). In secondary veins of maturing leaf 1, the basal ends of secondary veins showed delayed formation of wall ingrowths when compared to the apical ends (Fig. 1A). Notably, wall ingrowth deposition accumulated more rapidly in the juvenile foliar tissues (leaf 1) compared to leaf 7 which formed later during growth. Progression of wall ingrowth development from a score of below 3 to over 7 occurred across a nine-day window in leaf 1, but over 2 weeks in leaf 7. Together, these results suggest that the development of wall ingrowths in foliar tissues follows an underlying basipetal pattern but with temporal differences present between leaves of different developmental status. Furthermore, the analysis demonstrates that wall ingrowth deposition in PP TCs occurs well after structural development of individual minor veins.

### Distribution and morphology of PP TCs in minor veins of mature leaves

While wall ingrowth deposition in PP TCs has been studied by electron microscopy, the distribution of PP TCs relative to other phloem cells in individual veins remains unclear. Assessing the relative location of PP TCs in minor veins was investigated by collecting confocal optical stacks (Fig. 2A) and generating orthogonal reconstructions that were cross sections (Fig. 2B). Typically, a mature minor vein possessed two rows of PP TCs which were both located on the abaxial side of the phloem within the vascular bundle (Fig. 2B; Fig. 3A). However, a survey of 40 minor veins indicated that a relatively large percentage of minor veins had more than two PP TCs (around 45% as shown in Fig. 3C), and that these veins contained not only paired PP TCs localized on the abaxial side of the phloem but also additional PP TCs that were scattered throughout the vascular bundle (Fig. 3B, C). Moreover, analysis of minor vein size, calculated as the combined area of phloem and xylem tissues, suggested that the number of PP TCs in a minor was not correlated with vein size (Fig. 3C). A survey on 156 individual minor veins was conducted to further clarify relative location of PP TCs in minor veins. Counting the PP TCs that were directly adjacent to or just one cell away from the abaxial bundle sheath cells as the abaxial PP TCs, 59% of PP TCs located on the abaxial side of the phloem, while PP TCs on other sites (on the adaxial and middle sites) accounted for 41% (Fig. 3D). Moreover, abaxial PP TCs had the highest levels of wall ingrowth deposition compared to PP TCs positioned on the adaxial side of the phloem which had significantly lower levels (Fig. 3E; Supplementary Fig. 2). Together, wall ingrowth deposition showed a gradient in the phloem with abundance generally decreasing from the abaxial to the adaxial side of the phloem.

**Fig. 2.**
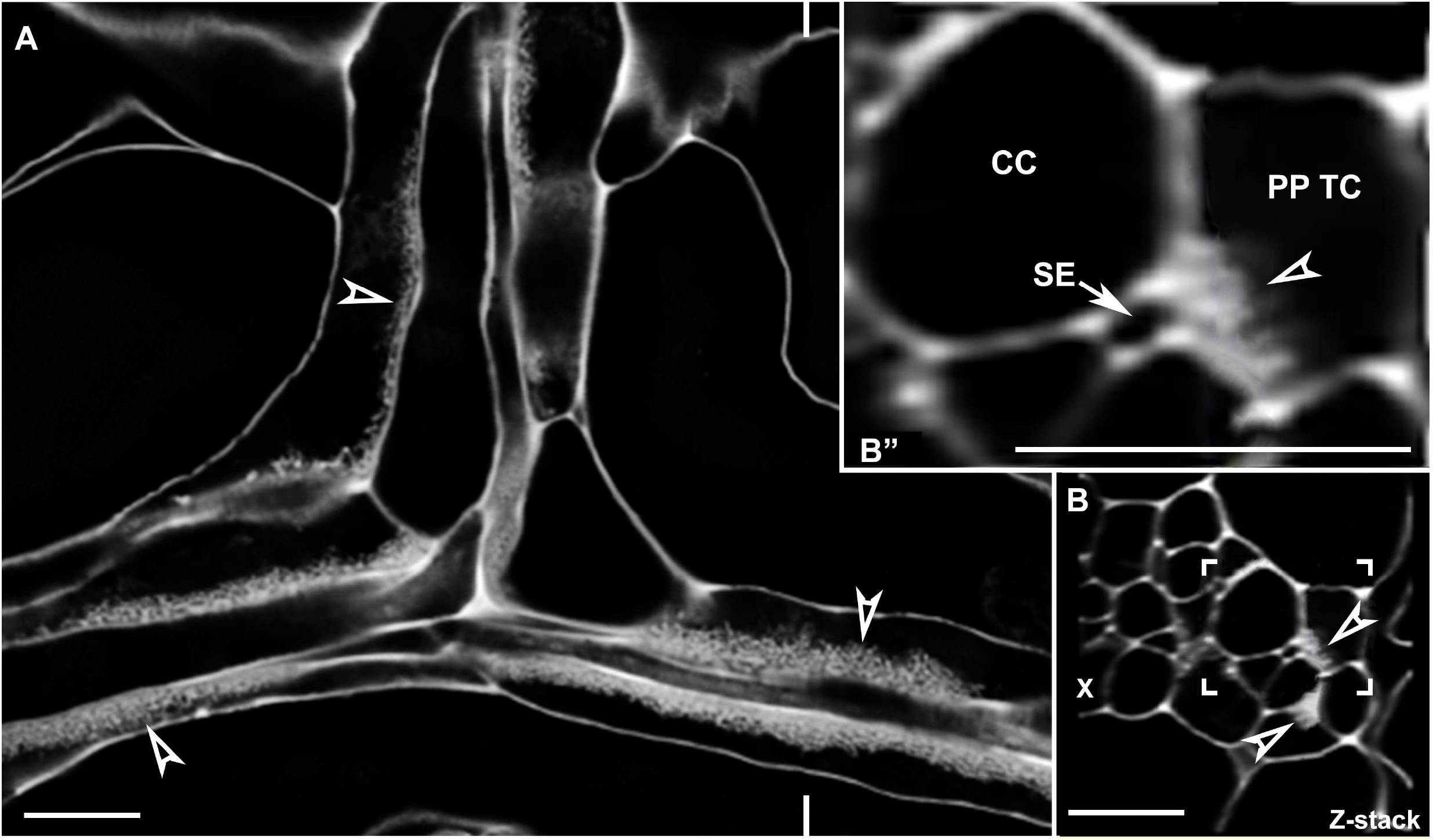
PP TC morphology and typical distribution in minor veins of mature leaves as assessed by confocal microscopy. **(A)** A single confocal section from an optical stack. In minor veins of a mature leaf, wall ingrowth deposition in PP TCs was abundant and highly polarized. Levels of wall ingrowth deposition varied in different PP TCs (arrowheads). **(B)** Orthogonal reconstruction of a confocal Z stack through a vein shown in **(A)** at the location marked by indent lines indicated that most wall ingrowth deposition occurred at sites adjacent to the PP-SE interface. **(B”)** Magnified image of the boxed region shown in **(B)**. Note that wall ingrowths (arrowhead) in the PP TC occurs primarily adjacent to the SE (arrow). Arrowheads indicate wall ingrowths; PP TC, phloem parenchyma transfer cell; SE, sieve element indicated by arrow; CC, companion cell; X, xylem. Pictures are representative images from different veins. Scale bars: 10 μm.

**Fig. 3.**
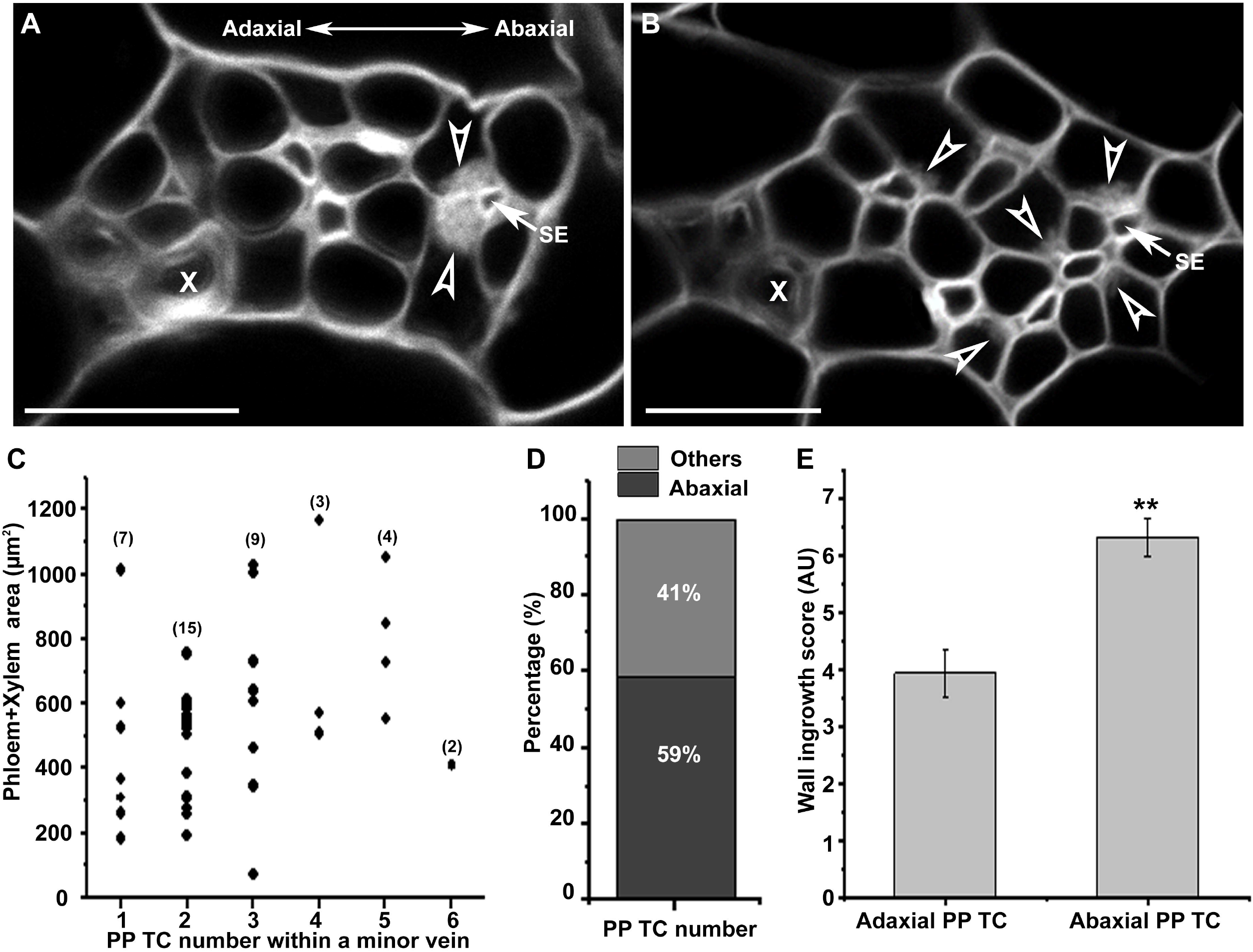
Wall ingrowth deposition was more abundant in abaxially-positioned PP TCs than adaxially positioned PP TCs. Confocal images were collected from mature leaf 7 from 5-week-old seedlings with inflorescences. **(A)** A typical leaf minor vein with two abaxially-positioned PP TCs containing wall ingrowths (arrowheads). **(B)** A leaf minor vein with multiple PP TCs (indicated by arrowheads denoting wall ingrowth deposition) positioned at different sites throughout the phloem. **(C)** Counts of the number of PP TCs per size of vascular bundle (phloem + xylem area) derived from cross sections of 40 minor veins. No correlation was seen between the number of PP TCs and the combined area of xylem and phloem cells, indicating that vein size does not control PP TC development. Numbers in brackets indicate the replicates from each minor vein class. **(D)** Proportions of PP TCs located on different sites of the phloem in leaf minor veins. From 339 PP TCs present in 156 minor veins, 200 PP TCs were abaxially-positioned PP TCs (defined as directly adjacent to or just one cell away from the abaxial bundle sheath cells) while 139 PP TCs were located in the middle and adaxial sites of the phloem. **(E)** Wall ingrowth scores were higher in abaxially-compared to adaxially-positioned PP TCs. Wall ingrowth scores were determined in eight minor veins (from three leaves) with multiple PP TCs, with the comparison made between the most adaxially- and most abaxially-positioned PP TC cells. Arrowheads indicate wall ingrowth deposition; PP TC, phloem parenchyma transfer cell; SE, sieve element (indicated by an arrow); X, xylem. Asterisks in **(E)** indicate significant differences (Student’s t-test, *P* < 0.01). Samples for (A-D) were Vibratome-cut cross sections whereas (E) used whole mount leaves. Scale bars: 10 μm.

### Developmental and spatial progression of wall ingrowth deposition in leaf minor veins

As shown in Fig. 3B, in mature minor veins, early-stage wall ingrowth deposition occurred almost exclusively at the PP TC/SE interface, suggesting a possible association between the localised deposition of wall ingrowths in PP TCs and adjacent SEs. To investigate this potential association, confocal images of cross sections of maturing leaf 7 from 4- to 5-week-old seedlings were surveyed. Following the scoring system introduced by Nguyen et al. (2017), five categories of wall ingrowth deposition were defined. Class I PP cells showed no discernible wall ingrowths (Fig. 4A) while in Class II, nascent wall ingrowths emerged at the PP TC/SE interface as evidenced by the bristle-like structure along the uniform primary cell wall (Fig. 4B). In Class III, wall ingrowths in PP TCs were more obvious and covered the entire PP TC/SE interface and sometimes covered a small section of the adjacent PP TC/CC interface (Fig. 4C), while in Class IV, wall ingrowths were more abundant and not only covered the entire PP TC-SE interface but also covered a considerable proportion of the adjacent PP TC/CC interface (Fig. 4D). In Class V, extensive wall ingrowths were abundant adjacent to SEs and had also spread across most of the PP TC/CC interface (Fig. 4E). These observations indicated that wall ingrowths were first deposited adjacent to SEs but then gradually extended to cover adjacent PP TC/CC interfaces, as indicated in the schematic figure (Fig. 4F).

**Fig. 4.**
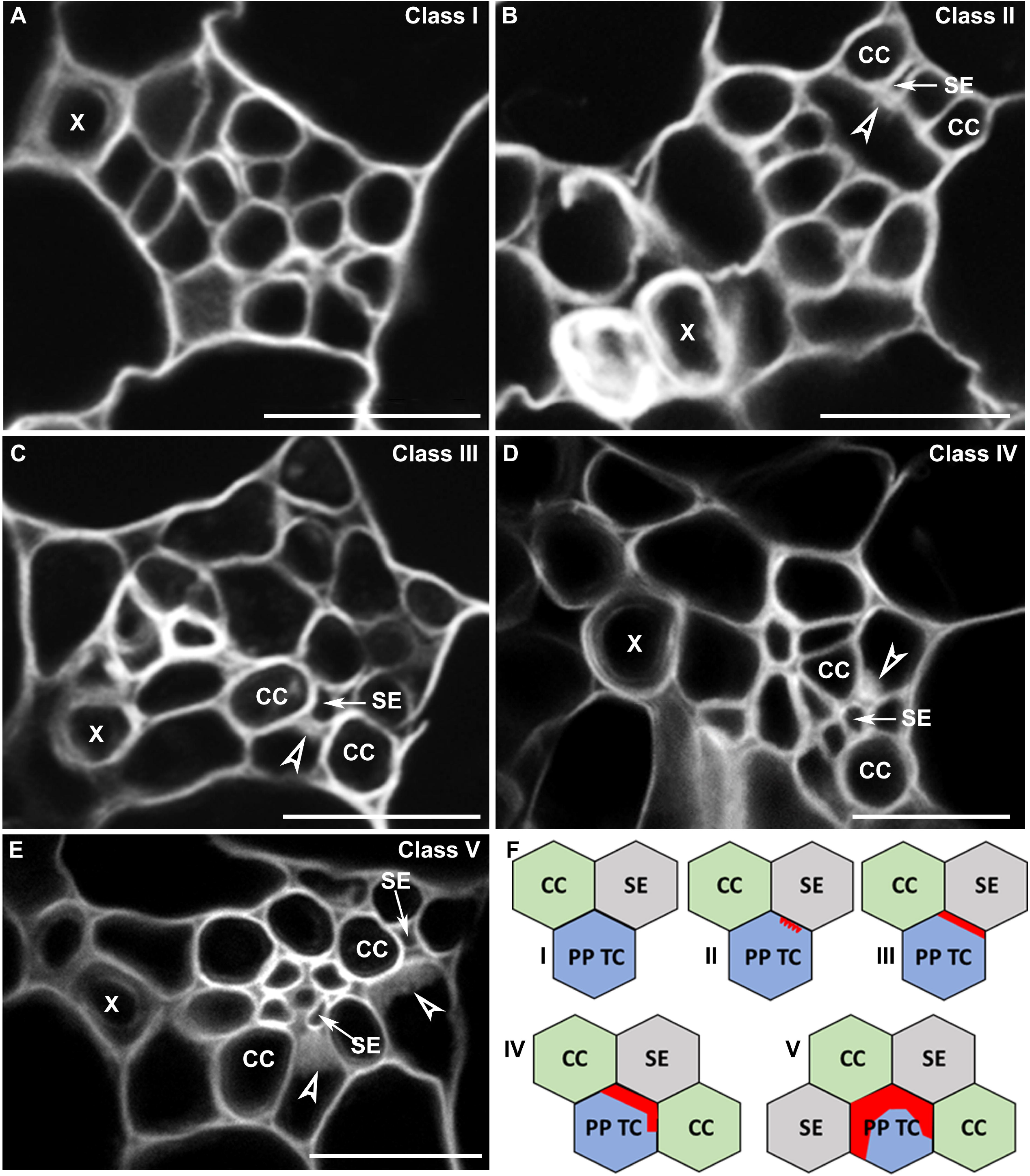
Development of wall ingrowth deposition in PP TCs in relation to neighbouring SEs and CCs. Cross section images were collected from Vibratome-cut cross sections through maturing leaf 7 of 4- to 5-week-old Col-0 seedlings and the extent of wall ingrowth deposition in PP TCs grouped into five representative classes. **(A)** Class I – no discernible wall ingrowths; **(B)** Class II – nascent wall ingrowth deposition positioned at the PP TC/SE interface; **(C)** Class III – substantial levels of wall ingrowth deposition entirely covering the PP TC/SE interface; (D) Class IV – extensive levels of wall ingrowth deposition cover the entire PP TC/SE interface and a small portion of the PP TC/CC interface; **(E)** Class V – massive levels of wall ingrowth deposition cover the entire PP TC/SE interface and considerable portions of neighbouring PP TC/CC interfaces. **(F)** Schematic figures illustrating the five representative classes of the extent of wall ingrowth deposition in PP TCs. Arrowheads indicate wall ingrowth deposition; arrows indicate sieve elements (SE); Red blocks in **(F)** indicate areas of wall ingrowth deposition as defined by Class I to Class V; PP TC, phloem parenchyma transfer cell; CC, companion cell; X, xylem. Images are representative images from three or more independent samples. Scale bars: 10 μm.

### Vascular structure and relative localization of wall ingrowths in mature minor veins in Arabidopsis leaves

TCs develop wall ingrowths to facilitate enhanced rates of *trans*-membrane transport of solutes (Pate and Gunning, 1972). In mature PP TCs, wall ingrowths cover the interface adjacent to the SE/CC complex (Haritatos et al., 2000; Amiard et al., 2007; Nguyen et al., 2017), suggesting existence of solute transport at the joint interface. However, the precise relationships between wall ingrowths in the PP TCs relative to SEs and CCs are poorly understood. Consequently, the function of PP TCs and their potential interactions with other phloem cells have not been fully elucidated Thus, to further understand the positioning of wall ingrowths relative to other cell types, in particular CCs, we used the transgenic line *pAtSUC2::AtSTP9-GFP (tmSTP9)*. The AtSTP9-GFP fusion expressed in this line is anchored to the CC plasma membrane and is a non-mobile CC marker (Stadler et al., 2005). Furthermore, activation of the *AtSUC2* promoter indicates functional maturation of minor veins (Wright et al., 2003). Thus, the expression of AtSTP9-GFP under control of the *AtSUC2* promoter not only reveals the identification of CCs but also provides a marker for the functional maturation of minor veins. By staining cross sections of minor veins from this line with calcofluor white to reveal wall ingrowths, it was possible to unambiguously identify CCs and PP TCs to enable the quantification of these cell types within vascular bundles.

Consistent with previous observations made by transmission electron microscopy (Haritatos et al., 2000; Amiard et al., 2007), our fluorescence observations confirm that wall ingrowth deposition in PP TCs in mature leaves occurs only at the interface of the cell wall adjacent to SEs and CCs (Fig. 5A), supporting the role of PP TCs in phloem loading (Chen et al., 2014). However, as shown in Fig. 5A, not all CCs in minor veins lie adjacent to PP TCs, suggesting the phloem loading path in leaf minor veins could be complex. From observations of 145 leaf minor veins, and counting cell types in minor veins in mature transition leaves 6, 7, and 8, the relative numbers of CCs, SEs and PP TCs are about 5, 4 and 2, respectively, in each minor vein (Fig. 5B).

**Fig. 5.**
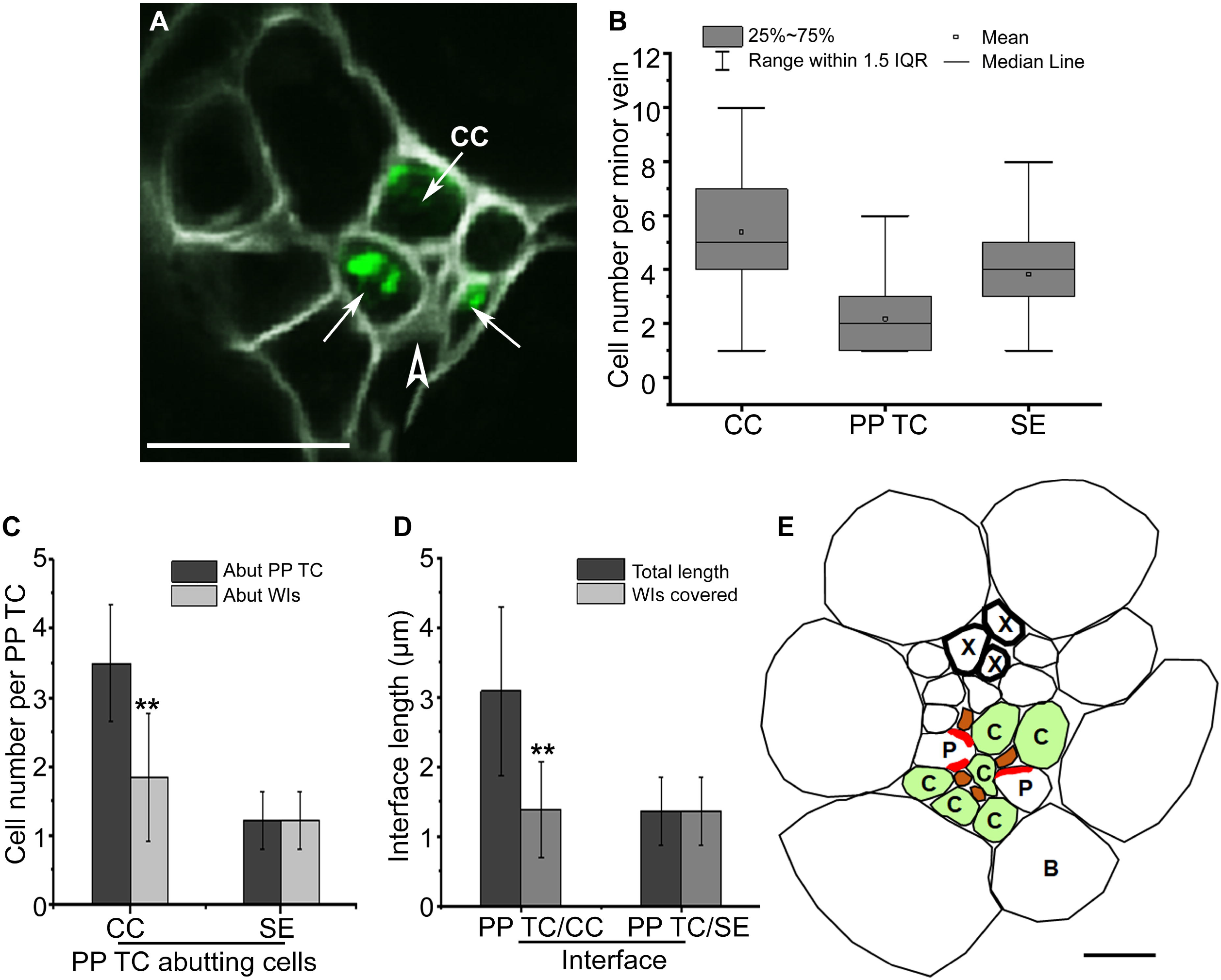
Minor vein structure and localization of wall ingrowth deposition in leaf vascular tissue in Arabidopsis. **(A)** Confocal imaging of Vibratome cross section through an unfixed minor vein from mature transition leaf 7 of the *pAtSUC2:AtSTP9-GFP* line expressing AtSTP9-GFP under control of the *pAtSUC2* promoter and localised to the plasma membrane of CCs (arrows). Sections were counter-stained with calcofluor white. Arrows indicate AtSTP9-GFP-expressing CCs, with the plasma membrane plasmolysed as a consequence of fixation. The arrowhead indicates wall ingrowths in the PP TC. **(B)** Phloem cell numbers in each minor vein of mature leaves. The average cell numbers were obtained from cross sections of 145 leaf minor veins. **(C)** Numbers of CCs and SEs that abut a PP TC (n = 36 PP TCs from 26 independent minor veins) (dark grey) and numbers of CCs and SEs around a PP TC that abut wall ingrowths (light grey). **(D)** Lengths of PP TC/CC (n = 160) and PP TC/SE (n = 84) interfaces, and lengths of the PP TC/CC and the PP TC/SE interfaces containing wall ingrowths. **(E)** Schematic diagram of vascular components in a minor vein of mature Arabidopsis leaves. X, Xylem; P, PP TC; C, CC; B, Bundle sheath. Red blocks in PP TC in **(E)** represent wall ingrowths. Asterisks in **(C)** and **(D)** indicate significant difference by the Student’s t-test (** *P* < 0.01). Scale bars: 10 μm.

The relative proportions and locations of the different cell types were also quantified. On average, one PP TC abuts approximately 3.5 CCs and 1.2 SEs, but only 55% of the CCs lie adjacent to wall ingrowths (Fig. 5C, E). In contrast, SEs that abut PP TCs all lie adjacent to wall ingrowths (Fig. 5C, E). Furthermore, for those PP TC/CC interfaces in which wall ingrowths are present, only 50% of the interfaces were covered by wall ingrowths whereas the entire SE interface of the PP TCs were covered with wall ingrowths (Fig. 5D, E). Interestingly, as shown in Fig. 5D, the average length of wall ingrowth along PP TC/CC interfaces was almost equal to that of the PP TC/SE interface, being 1.39 μm and 1.36 μm, respectively. These observations suggest that the association between wall ingrowth deposition in mature PP TCs abutting SEs is stronger than that with CCs. Together, these observations indicate the impact of abutting SEs on wall ingrowth formation on PP TCs is functionally more significant than that of the abutting CCs in leaf minor veins in Arabidopsis.

### Development of PP and PP TCs in leaf minor veins in Arabidopsis

AtSWEET11 is a sucrose transporter localized to the plasma membrane of PP TCs where it functions to export sucrose from PP TCs into the apoplasm adjacent to the SE/CC complex in leaf minor veins in Arabidopsis (Chen et al., 2012; Cayla et al., 2019). However, a recent study using single-cell sequencing (scRNA-seq) analysis demonstrated that leaf PP cells expressing *AtSWEET11* can be classified into two distinct clusters. Genes involved in callose deposition and cell wall thickening are over presented in one PP cluster (PP1), whereas genes involved in photosynthesis are enriched in the second cluster (PP2) (Kim et al., 2021). This data implies the existence of two distinct populations of PP cells in Arabidopsis leaves.

Based on these observations, the hypothesis arises that PP TCs with wall ingrowths belong to the PP1 cluster, while PP cells with no wall ingrowths belong to the PP2 cluster. Previous studies have suggested that PP cells expressing *AtSWEET11* and *AtSWEET12* have wall ingrowths (Chen et al., 2012; Cayla et al., 2019). However, whether all PP cells in leaf minor veins have wall ingrowths, thus defining them as PP TCs, remains unknown. To clarify this question, mature juvenile leaf 1 from 27-day-old seedlings of the *pAtSWEET11::AtSWEET11:GFP* transgenic line were observed by confocal microscopy. However, PP TCs in Arabidopsis are located in vascular bundles, which makes imaging of PP TCs in whole leaves is particularly challenging. Therefore, peeling away the abaxial epidermis as described by Cayla et al. (2019), followed by fixation and ClearSee treatments (Kurihara et al., 2015) were applied. Cleared samples were then stained with calcofluor white to observe cell walls. In Fig. 6, a minor vein containing two cell files expressing AtSWEET11-GFP, thus identified as PP cells, is presented. Orthogonal reconstructions showed two PP cells, labelled 1 and 2 (Fig. 6A-C) with different characteristics. For example, PP cell 1 was adjacent to a SE (asterisk) but PP cell 2 was not. More importantly, PP cell 1 had *trans*-differentiated into a PP TC as evidenced by abundant wall ingrowth deposition along its interface adjacent to the SE (Fig. 6E). In contrast, PP cell 2 had no detectable levels of wall ingrowth deposition (Fig. 6G), indicating that not all AtSWEET11-GFP-expressing PP cells in the Arabidopsis leaf minor vein had *trans*-differentiated to become PP TCs. However, not all observed minor veins showed this pattern, with analysis of 20 minor veins from eight different leaves revealing that only five minor veins contained PP cells expressing AtSWEET11-GFP but not depositing wall ingrowths to become PP TCs. In most cases, multiple PP cells within a minor vein all deposited wall ingrowths along the interface adjacent to the SE, as shown in Supplementary Fig. 3. Together, these observations suggest that minor veins have two sub-types of PP cells, one without discernible wall ingrowth deposition, and the other having *trans*-differentiated into PP TCs as a consequence of wall ingrowth deposition. In addition, these results also confirm that adjacent SEs appear to be necessary for the wall ingrowth deposition in PP TCs.

**Fig. 6.**
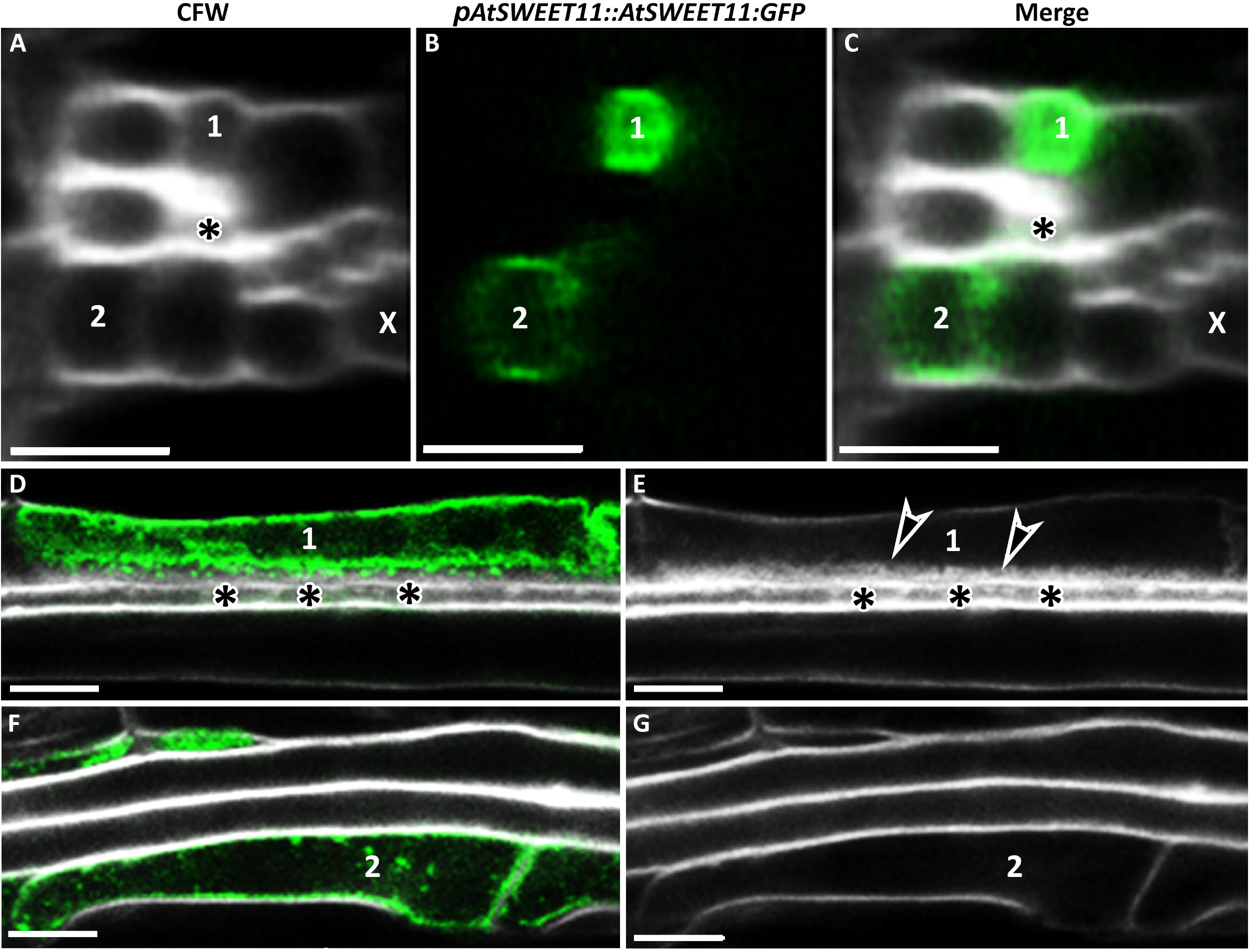
Adjacent SEs are required for wall ingrowth deposition in PP TCs in minor veins of Arabidopsis. Leaf 1 from 27-day-old seedlings of the transgenic line *pAtSWEET11::AtSWEET11-GFP* was fixed and cleared using ClearSee, stained with calcofluor white (CFW) and imaged by confocal microscopy. **(A-C)** Cross-sections were generated from confocal optical stacks using orthogonal sectioning. **(A)** Cross section of a minor vein in a mature juvenile leaf stained with calcofluor white and showing two PP cells (labelled 1 and 2) and an adjacent SE (asterisk). **(B)** AtSWEET11-GFP fluorescence in PP cells 1 and 2. **(C)** Merged image of A and B, indicating the presence of two PP cells showing AtSWEET11-GFP expression. PP cell 1 was adjacent to a SE (asterisk) and showed strong AtSWEETll-GFP fluorescence, whereas PP cell 2 showed weaker fluorescence and was not immediately adjacent to a SE. **(D, E)** PP cell 1 expressing AtSWEET11-GFP **(D)** showed abundant wall ingrowth deposition (arrowheads in **E)** on the interface adjacent to a SE (asterisk). **(F, G)** PP cell 2 had no detectable wall ingrowth deposition (G), but expressed AtSWEET11-GFP **(F)**. X, Xylem. Arrowheads indicated wall ingrowth deposition. Asterisks indicate SE. Results shown here are representative of four minor veins. Scale bars: 5 μm.

### Accumulation of AtSWEET11-GFP in PP and PP TCs

Since wall ingrowth deposition in TCs results in increased plasma membrane surface area and AtSWEET11 is localised to the plasma membrane in PP TCs, it is reasonable to assume that AtSWEET11-GFP should be abundant in regions of wall ingrowth deposition in these cell types in minor veins. To investigate this hypothesis, fluorescence intensity of AtSWEET11-GFP was determined in foliar minor veins using *pAtSWEET11::AtSWEET11-GFP* transgenic plants. Since the processing of tissue for Vibratome cross sectioning induces plasmolysis (Fig. 5A), images of fixed and ClearSee-treated plants were collected as optical stacks by confocal microscopy, and then images were orthogonally reconstructed to provide cross sections. In Arabidopsis leaves, the phloem is located beneath the epidermis and bundle sheath cells. Consequently, images of adaxially-positioned phloem cells collected by confocal microscopy will be darker compared to phloem cells located at more abaxially-positioned sites, since light scattering and absorption by cells that occur in the light path weaken excitation light intensity and reduces emissions. To address this limitation, calcofluor white fluorescence of general cell wall staining was used as an internal standard to calibrate AtSWEET11-GFP fluorescence intensities of different PP cells (Fig. 7A, B).

**Fig. 7.**
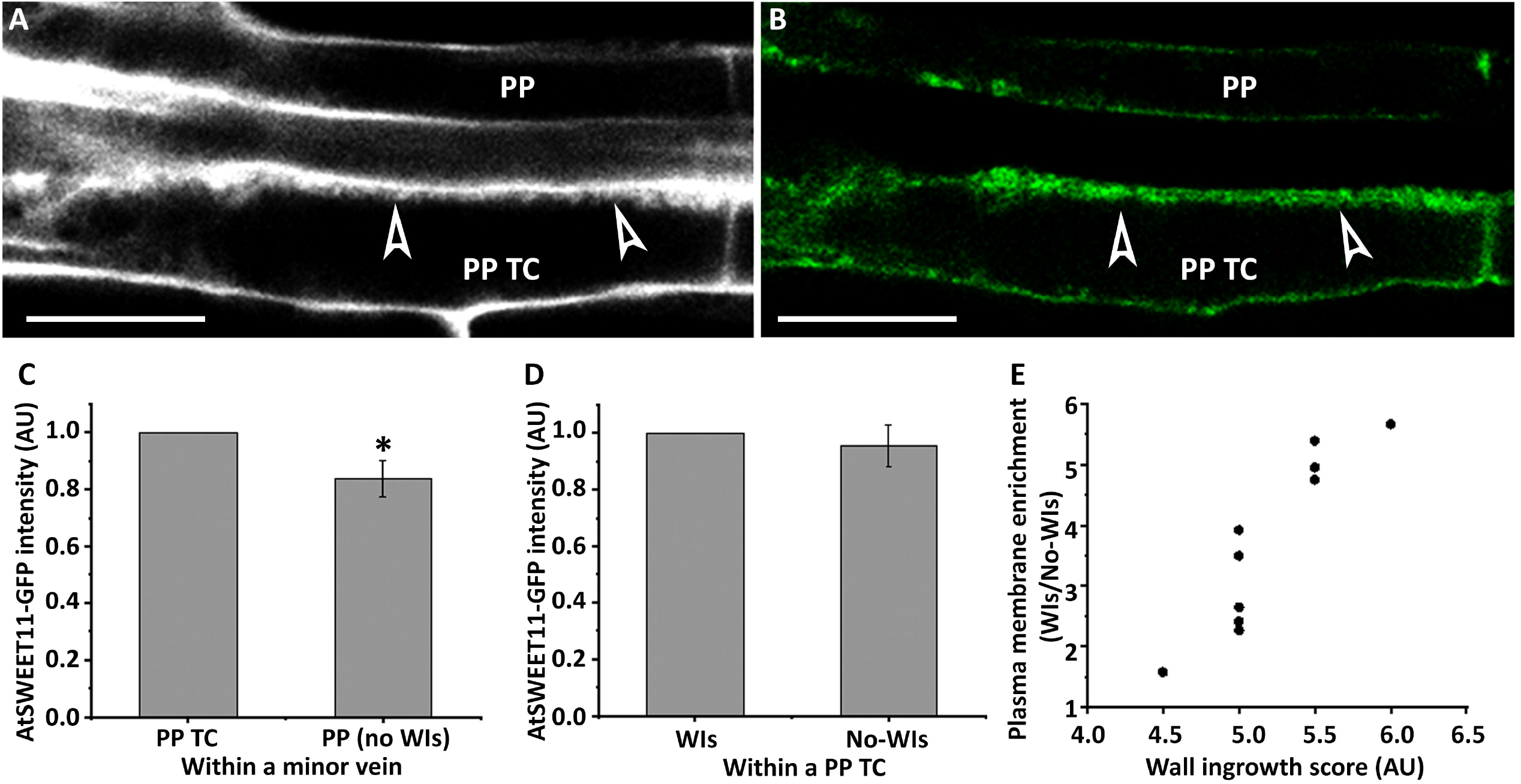
Wall ingrowth deposition positively associates with AtSWEET11 accumulation in PP cells. Leaf 1 from 27-day-old seedlings were fixed in 4% (w/v) formaldehyde, cleared using ClearSee solution, and stained with calcofluor white. **(A)** Cell wall labelling using calcofluor white. Arrowheads indicate wall ingrowth deposition in the PP TC, but no deposition in the PP cell. **(B)** AtSWEET11-GFP fluorescence of the same region shown in A. **(C)** Relative fluorescence intensity of AtSWEET11-GFP in PP TCs and PP cells in minor veins. **(D)** Relative fluorescence intensities of AtSWEET11-GFP in PP TCs where wall ingrowth (WI) deposition had occurred or not. Four pairs of PP TC and PP cell from four different leaf samples were compared and analysed to generate **(C)** and **(D)**. **(E)** Plasma membrane accumulation, as measured by the ratio of AtSWEET11-GFP fluorescence on the side of the cell with wall ingrowths against the opposite side of the cell, as a function of wall ingrowth deposition. Data obtained from ten PP TCs in eight different leaves. Asterisk in **(C)** indicates significant difference by Student’s t-test (\**P* < 0.05). Pictures are representative images from four independent samples. Scale bars: 10 μm.

In minor veins, the AtSWEET11-GFP signal was stronger in PP TCs compared to PP in which no wall ingrowth deposition had occurred (Fig. 7C). This analysis supports the qualitative observations made in Fig. 6. Statistical analysis indicated that the fluorescence intensity of AtSWEETll-GFP in PP cells was about 84% of the signal intensity detected in PP TCs (Fig. 7C). However, fluorescence intensity of the AtSWEET11-GFP signal showed no distinct difference between sites with wall ingrowths and those without (Fig. 7D), indicating AtSWEET11 is relatively evenly distributed along the plasma membrane in PP TCs. Since SWEET11-GFP is plasma membrane-localized and SWEET11-GFP does not localise preferentially to plasma membrane surrounding wall ingrowths (Fig. 7D), plasma membrane area per unit length at the site with wall ingrowths could be quantified by measuring the accumulation of SWEET11-GFP fluorescence in PP TCs. As shown in Fig. 7E, wall ingrowth deposition in PP TCs resulted in localised plasma membrane enrichment. For instance, compared to sites with no wall ingrowths, plasma membrane area at sites with a wall ingrowth score of five was about three-fold higher than sites with no wall ingrowths (Fig. 7E). Moreover, the sites with higher levels of wall ingrowths, as indicated by ingrowth scores, had larger plasma membrane areas (Fig. 7E). Consequently, by achieving an enlarged plasma membrane surface area, wall ingrowth deposition enables higher local densities of AtSWEET11 compared to other areas in PP TCs.

## Discussion

### Onset of wall ingrowth deposition in PP TCs is coupled with functional maturation in Arabidopsis leaf minor veins

In Arabidopsis leaves, the structural maturation of veins proceeds basipetally with the lower order veins (midrib and secondary veins) developing first (Kang et al., 2007). Once the formation of the midrib is finalized, the first loop of secondary veins develops outwards from the midrib and acropetally until a complete loop is formed. Afterwards, the second loop starts to form in a bidirectional manner, with development initiated from both the first loop as well as the midrib (Scarpella et al., 2006; Scarpella et al., 2010), with the same sequence happening for the formation of the third loop and subsequent loops. The results presented in this study illustrate the spatial and temporal developmental progression of PP TCs in Arabidopsis. The onset of PP TC development in veins starts at a relatively later stage of leaf development when vascular patterning is close to completion (Fig. 1). In other words, wall ingrowths do not form until the vein is structurally mature, indicating that the formation of wall ingrowths in PP TCs is not coupled with the structural maturation of the vein. This conclusion is further supported by the observation that the higher-order minor veins, such as quaternary veins that are formed last, will develop similar (or even more extensive) levels of wall ingrowth compared to the lower order, secondary and tertiary veins that are formed earlier in leaf development (Fig. 1). Coinciding with the development of wall ingrowths, the sink-to-source transition (functional maturation process) in foliar tissues also proceeds from the tip to the base (Wright et al., 2003; Wei et al., 2020). Moreover, consistent with the observation in a previous study (Nguyen et al., 2017), the heteroblastic development of PP TCs was observed with distinct differences presented between leaf 1 and leaf 7 (Fig. 1). Compared to leaf 7, the basipetal gradient of the sink-to-source transition was much less obvious in leaf 1 in Arabidopsis (Supplementary Fig. 4), which is consistent with their temporary difference in wall ingrowth deposition. Together, these observations showed a tight correlation between wall ingrowth deposition and functional maturation process of the phloem, which not only provides further evidence for the positive correlation between wall ingrowth deposition and phloem loading activity in PP TCs, but also offers new insights into the underlying mechanism for the heteroblastic development of PP TCs in Arabidopsis.

### Arabidopsis leaf minor veins contain two sub-types of PP cells

Several recent studies have demonstrated the structural and functional complexity of the phloem tissue in plants. In grasses such as maize and rice, two types of SEs have been identified, one thin-walled and symplasmically connected to CCs but no other cell type, while the other is thick-walled and having abundant plasmodesmatal connections to vascular parenchyma cells (McCubbin and Braun, 2021). Furthermore, in maize leaves, abaxially-positioned bundle sheath cells specifically express three SWEET sucrose transporters (SWEET13a, b and c) which are all involved in phloem loading, while adaxial bundle sheath do not express these transporters, indicating at least two different bundle sheath populations in maize leaves (Bezrutczyk et al., 2021). In Arabidopsis and tobacco leaf minor veins, two types of CCs have also been identified, one expressing *FT* and the other not (Chen et al., 2018). A recent single-cell RNA expression study also demonstrated the complexity of phloem with Kim et al., (2021) reporting that PP cells can be transcriptionally divided into two sub-clusters, a PP1 cluster enriched in cell wall genes and genes involved in transmembrane transport, and a PP2 cluster enriched in photosynthesis genes (Kim et al., 2021).

These observations demonstrate that phloem structure in leaves is complex and that phloem cell population including SEs, CCs and bundle sheath cells are functionally diverse. The results reported in this study expand this complexity to include PP cells in Arabidopsis minor veins and suggests the presence of two sub-types of PP cells - one sub-type adjacent to SEs and containing wall ingrowths, thus defining them PP TCs, while other PP cells in the same minor vein do not develop wall ingrowths and were not adjacent to SEs (Fig. 6). Compared to PP cells, PP TCs not only contained wall ingrowths but also more strongly expressed *AtSWEET11* (Fig. 6; Fig. 7), indicating morphological and physiological differences between these two subtypes of PP cells. Together, these results strongly suggest the presence of two sub-types of PP cells in leaf minor veins in Arabidopsis. In this context, it would be interesting to investigate whether the PP1 cluster identified by Kim et al. (2021) corresponds to PP TCs and thus the PP2 cluster corresponds to PP cells.

### Phloem loading might preferentially occur via abaxial PP TCs in Arabidopsis leaves

Results from this and a previous study (Wei et al., 2020) demonstrate that wall ingrowth deposition is positively correlated with phloem loading activity in PP TCs. Therefore, wall ingrowth deposition could be regarded as a trait, similar to root growth, that indicates phloem loading activity in Arabidopsis. In most mature minor veins, PP TCs mostly develop on the abaxial-side of the phloem (Fig. 2B; Fig. 4B-E; Fig. 5A). Moreover, PP TCs are more active in phloem loading as they possess higher levels of AtSWEET11 compared to non-*trans*-differentiated PP cells that have no wall ingrowths (Fig. 7). Furthermore, in minor veins that have multiple PP TCs, wall ingrowth deposition levels also show a distinct gradient so that PP TCs on the abaxial-side of the phloem accumulated more abundant wall ingrowths in comparison with the PP TCs on the adaxial-side (Fig. 5). Thus, one might hypothesise that the phloem loading activity in PP TCs that are localized on the abaxial-side of the phloem might be higher than those on the adaxial side. Evidence from scRNA-seq analysis of maize leaves established that sucrose uptake into phloem is mainly via abaxial bundle sheath cells (Bezrutczyk et al., 2021). Consistent with this observation, the path of sucrose export in maize minor veins is through the abaxial-positioned thin-walled SEs (reviewed by McCubbin and Braun, 2021). These observations demonstrate that phloem loading activity is asymmetric in maize minor vein, which supports the hypothesis developed here regarding the different functions for abaxial and adaxial PP TCs in phloem loading in Arabidopsis.

### Wall ingrowth deposition in PP TCs is tightly correlated to the adjacent SE in leaf minor veins in Arabidopsis

A body of evidence, including observations in this study, indicates that wall ingrowth deposition in PP TCs in Arabidopsis polarize to the interface adjacent to the SE/CC complex (Haritatos et al., 2000; Amiard et al., 2007; Adams et al., 2014). As shown in Fig. 4, in Arabidopsis minor veins, wall ingrowths first occurred at the PP TC/SE interface and then gradually extended across adjacent PP TC/CC interface, implying a potential impact of SEs on the initiation of wall ingrowths in PP TCs. Moreover, all the PP TC/SE interfaces were totally covered by wall ingrowth deposition, while less than half of the total length of the PP TC/CC interfaces were covered similarly (Fig. 5). This result complicates the assumption that wall ingrowths in PP TCs are developed by the plant mainly to facilitate solute exchange between PP TCs and CCs. In classical phloem loading models, neither the symplastic nor the apoplastic loading process has described any PP TC/SE interaction in the collection veins. Instead, the typical loading route proposes that sucrose exported from PP cells is actively taken up into CCs via SUC2/SUT1, and then delivered symplasmically into SEs for long-distance transport (Turgeon and Wolf, 2009; Schepper et al., 2013; Chen et al., 2014). Thus, the polarised distribution of wall ingrowths, predominantly correlating spatially with SEs rather than CCs, seems peculiar. Furthermore, none of the minor vein cross section images collected in this study showed any PP TC with wall ingrowths that were not adjacent to SEs. However, observations in the *pAtSWEET11:AtSWEET11:GFP* transgenic plants indicated that a population of PP cells did not form discernible wall ingrowths and these PP cells were not adjacent to the SEs (Fig. 6). Together, the data reported here indicate that the *trans*-differentiation of PP cells into PP TCs is tightly associated with adjacent SEs in Arabidopsis.

### Reconsidering the role of wall ingrowth deposition in phloem loading in Arabidopsis leaves

Wall ingrowth deposition in PP TCs promotes phloem loading by enlarging plasma membrane surface area of the exporting PP TCs (Fig. 7). However, our previous study showed that wall ingrowth formation is induced by phloem loading, rather than wall ingrowth deposition itself being a prerequisite required for phloem loading (Wei et al. 2020). Wall ingrowths in PP TCs are unlikely to make a significant contribution to sucrose loading until they reach high levels (scores of five or higher) because until that stage, they do not greatly enhance plasma membrane area (Fig. 7) (see also Rae et al. 2021. In addition, mature transition and adult leaves show low levels of wall ingrowth deposition towards the base of the leaf with scores of less than 4 (Nguyen et al., 2017; Fig. 1) which seems to contradict a role for wall ingrowths in facilitating *trans*-membrane transport of photo-assimilates in these cells. Furthermore, as discussed in this study, wall ingrowth deposition preferentially covers the PP TC/SE interface rather than the CC/SE interface (Fig. 3, 4, and 5). However, the PP/CC interface is proposed to be the site where phloem loading of sucrose occurs (Chen et al., 2014). Together, these observations indicate that wall ingrowth deposition in PP TCs may have additional roles rather than solely functioning in facilitating sucrose transport. Interestingly, the steady-state pattern of hydrostatic pressure distribution in the leaf phloem network, which is largely determined by leaf shape, coincides with the heteroblastic pattern of wall ingrowth deposition observed in Arabidopsis (reviewed by Wei et al., 2021). The hydrostatic pressure in the leaf is caused by solute uptake in collection phloem (Gould et al., 2005). Similar to the distribution of wall ingrowth deposition in Arabidopsis leaves, the distribution of hydrostatic pressure in narrower/longer leaves shows a basipetal gradient, whereas it is relatively uniform in wider/shorter leaves (Sakurai and Miklavcic, 2021). These similarities imply the existence of a possible association between wall ingrowth deposition and water transport in Arabidopsis leaves.

## Conclusions

We have demonstrated that wall ingrowth deposition enhances phloem loading activity by increased accumulation of the sucrose transporter AtSWEET11 in PP TCs. Moreover, results from this study refine the temporal and spatial development of PP TCs, which provides further evidence for the impact of phloem loading activity on wall ingrowth deposition in PP TCs (Wei et al., 2020). The asymmetric distribution patterns of wall ingrowth deposition across minor veins in Arabidopsis, in turn, imply that phloem loading activity in Arabidopsis leaves is heteroblastic, and might be mainly undertaken by abaxial PP TCs in minor veins. However, our observations demonstrate the influence of SEs on PP TC development is more dominant compared to that of CCs, implying potential interaction between PP TCs and SEs in phloem loading. Furthermore, results from this study also suggests that there might be two sub-types of PP cells exist in leaf minor veins in Arabidopsis that are distinguished by the presence of wall ingrowths.

## Supplementary data

The following supplementary data are available at JXB online.

*Supplementary Fig. 1*. PP TC morphology and distribution in the midrib of mature leaves.

*Supplementary Fig. 2*. Representative images of adaxially- and abaxially-positioned PP TCs in a mature leaf minor vein.

*Supplementary Fig. 3*. Representative image of a mature minor vein showing that all PP cells become PP TCs.

*Supplementary Fig. 4*. Functional maturation status in maturing leaves in Arabidopsis.

## Acknowledgements

We thank the following for seeds: ABRC (Col-0); Dr. Ruth Stadler *(pAtSUC2::AtSTP9-GFP, tmSTP9)*; Dr. Sylvie Dinant *(pAtSWEET11::AtSWEET11)*.

## Author contributions

XW, DC and DM devised the project; XW performed experimental analyses; XW, YH, DC and DM analysed the data. XW, DC and DM wrote the manuscript and all authors approved the final text.

## Data availability

All data supporting the findings of this study are available within the paper and within its supplementary data published online.

